# DNA metabarcoding reveals the threat of rapidly expanding barred owl populations to native wildlife in western North America

**DOI:** 10.1101/2022.04.19.488820

**Authors:** Nicholas F. Kryshak, Emily D. Fountain, Daniel F. Hofstadter, Brian P. Dotters, Kevin N. Roberts, Connor M. Wood, Kevin G. Kelly, Isabel F. Papraniku, Paige J. Kulzer, Amy K. Wray, H. Anu Kramer, John P. Dumbacher, John J. Keane, Paula A. Shaklee, R.J. Gutiérrez, M. Zachariah Peery

## Abstract

Invasive predators can have detrimental impacts on native species and biological communities through direct consumptive effects and indirect effects on trophic interactions. As an invasive, apex predator achieving high densities, barred owls (*Strix varia*) may pose a substantial threat to native wildlife in western North American forests. Studies of the trophic ecology of barred owls in their invasive range, however, have involved morphological examinations of prey remains with limited taxonomic resolution. We conducted DNA metabarcoding using intestinal samples collected from barred owls at the leading edge of their range expansion in northeastern California. Using customized primers, we screened the intestinal contents of 124 barred owls and detected a broad diet of 78 unique prey types (48 vertebrates and 30 invertebrates), including many previously undetected prey types. Mammals were the most consumed vertebrate class (frequency of occurrence = 65%), followed by amphibians (32%), birds (22%), and reptiles (19%). Diets differed regionally but were similar among ages and sexes and exhibited limited variation in response to local environmental conditions. Our work highlights the generalist predatory strategy of invasive barred owls, indicates that they will not serve as ecological replacements for the congeneric spotted owls (*S. occidentalis*) they displace, and identifies numerous native species potentially threatened by their range expansion. Expanding currently limited barred owl removals could benefit native species and wildlife communities in western North America. More broadly, DNA metabarcoding provides a powerful tool for conducting detailed assessments of species consumed by invasive predators, potentially incentivizing conservation actions and improving outcomes.

## 1. Introduction

Species’ distributions are increasingly shifting in response to habitat alteration, climate change, transportation, and other human activities (Hobbs, 2000; Parmesan, 2006; Dornelas et al., 2014). When species enter new environments, they can become invasive — triggering a host of novel ecological interactions between historically isolated species that can threaten native species (Urban et al., 2012). Invasive predators can have particularly detrimental effects on native species through direct consumptive effects and are a leading cause of species extinctions globally (David et al., 2017). The consumption of native species by invasive predators can also cause significant changes to biological communities by altering and disrupting trophic linkages (Sih et al., 2010). Accordingly, efforts to mitigate the effects of invasive predators can benefit substantially from a rigorous understanding of what prey species they consume.

The ongoing range expansion of eastern barred owls (*Strix varia*) poses substantial concerns for native species and biological communities in western North American forests. For more than a century, this apex predator has been expanding its range westward through Canadian forests and riparian corridors across the Great Plains (Livezey, 2009). It has more recently undergone rapid population increases and achieved high abundance in the Pacific Northwest and parts of California (Forsman et al., 2011; Yackulic et al., 2012; Wiens et al., 2011, 2014; Wood et al., 2020). Barred owls have become entirely sympatric with — and greatly outnumbers — their congener, the northern spotted owl (*S. occidentalis caurina*), in this region (Gutiérrez et al., 2007). More recently, barred owls have colonized the range of the California spotted owl (*S. o. occidentalis*) in the Sierra Nevada, California, which represents the leading edge of their range expansion (Dark et al., 1998; Wood et al., 2019). Invasive barred owls and native spotted owls have been previously shown to occupy similar forest habitats and dietary niches, and the more dominant barred owl displaces territorial spotted owls via both exploitative (direct) and interference (indirect) competition (Hamer et al., 2001; Wiens et al., 2014; Wood et al., 2021). Indeed, barred owls are now considered an existential threat to spotted owls, with formerly large populations of spotted owls functionally extirpated (Gutiérrez et al., 2007; Wiens et al., 2021). In response, experimental removals of barred owls have been conducted at local scales in the range of the northern spotted owl (Diller et al., 2014; Wiens et al., 2021), and region-wide experimental removals have effectively reduced the population of barred owls in the Sierra Nevada (Hofstadter et al., 2022).

Barred owls, as generalist predators capable of achieving exceptionally high densities (Hamer et al., 2007; Wiens et al., 2014), may have broader effects on native species and biological communities than merely impacting spotted owls (Holm et al., 2016). In the Pacific Northwest, morphological analyses of prey remains contained in stomachs and regurgitated pellets indicate that barred owls consume a broad range of vertebrate and invertebrate taxa, including several species considered sensitive in that region (Hamer et al., 2001; Wiens et al., 2014). However, morphological analyses of pellet and stomach contents can lead to the underrepresentation of prey taxa (Raczynski and Ruprecht 1974), particularly small- or soft-bodied organisms (see: Livezey, 2007). Thus, the full suite of prey consumed — and potentially impacted — by this invasive and abundant apex predator in western North American forests has not been characterized.

We used DNA metabarcoding applied to samples collected as a part of removal studies to (*i*) conduct a comprehensive assessment of prey species consumed by barred owls and (*ii*) characterize how extrinsic and intrinsic factors shape the diet of barred owls along the leading edge of their invasion in the Sierra Nevada and southern Klamath/eastern Cascade (henceforth: Klamath/Cascade) ecoregions of northeastern California. DNA metabarcoding of digestive contents offers a potentially valuable means of quantifying owl diet that provides high taxonomic resolution, even for rare, highly degraded, or visually absent prey items (Piñol et al., 2014). Importantly, metabarcoding has been found to increase the proportion of identified prey items compared to visual methods in a variety of taxa (Newmaster et al., 2013; Pompanon et al., 2012), but has not yet been used to study the trophic ecology of barred owls. Our findings highlight species at direct, exploitative risk from invasive barred populations, as well as species that may face indirect pressure from interference competition.

## 2. Materials and Methods

### 2.1. Sampling of Barred Owls

We sampled from barred owls collected as part of a larger removal experiment in northeastern California (see: Hofstadter et al., 2022 for further detail; Fig. 1). Individuals were collected from September through November 2018 in the Klamath/Cascade and April through November 2019 in both ecoregions, under state and federal collection permits (California Department of Fish and Wildlife Service SCP-002114, SCP-11963, and United States Fish and Wildlife permits MB24592D-0, MB53229B-0). Removal procedures were consistent with approved IACUC protocol A006106-A01. Upon collection individuals were stored at -20°C, and delivered to the University of California, Berkeley Museum of Vertebrate Zoology for cataloguing and sampling. We aged barred owls as either adult (≥ 3 years), subadult (1-2 years), or juvenile (0 years) based on plumage and a wider terminal band on after-hatch-year flight feathers (Chris Neri, *personal communication*). We determined sex during the preparation of study skins by observing gonads. Intestines were collected for dietary analyses using aseptic techniques. Following collection, intestines were stored in 96% ethanol at -20°C (see Table A1 for sample archiving). Only intestinal material was collected for DNA extraction because (*i*) this allowed for complete collection of largely homogenized material and (*ii*) this methodology preserved stomach contents for future morphological comparisons.

**Fig 1.**
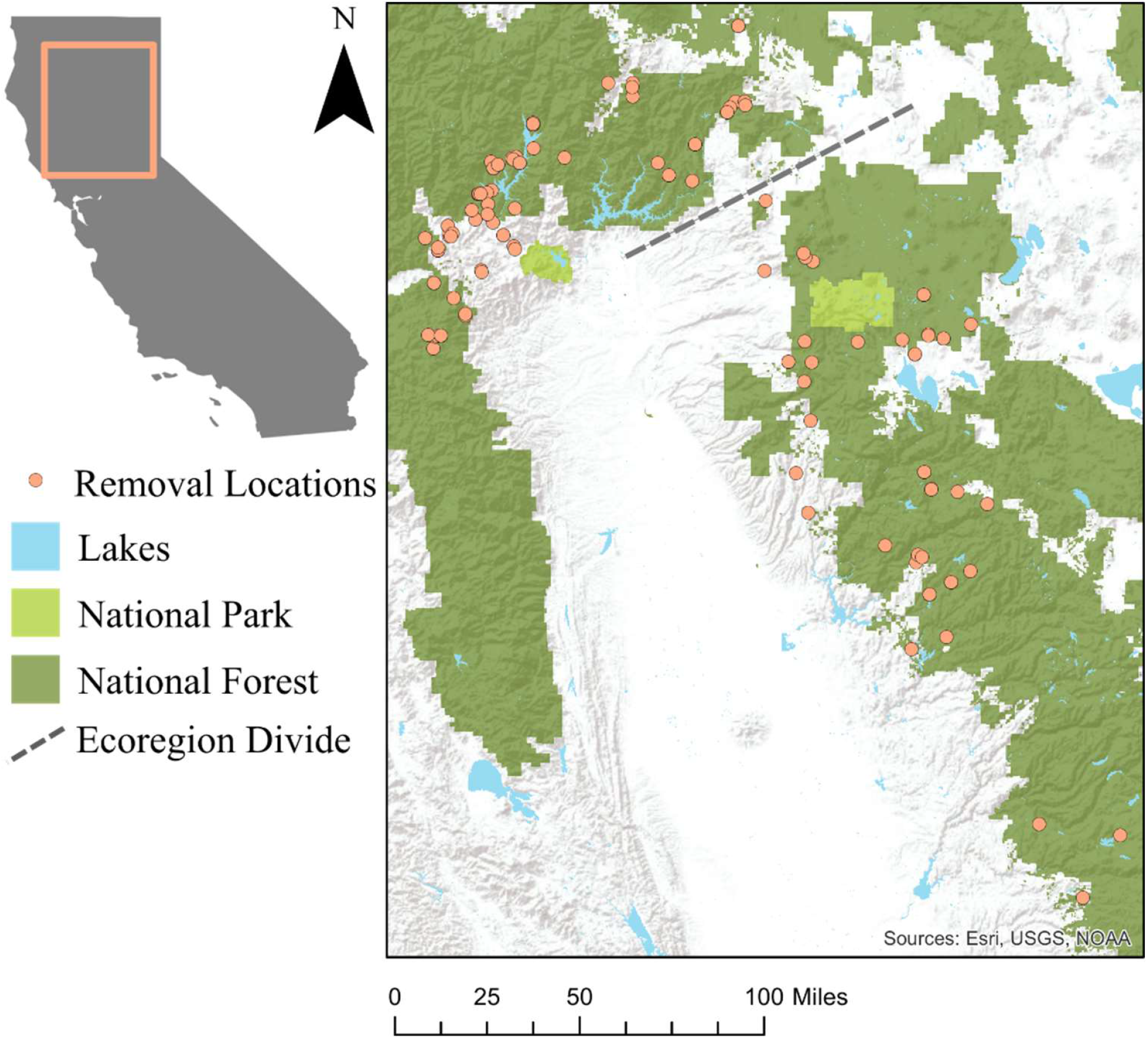
Barred owl removal locations in California, USA, in the Klamath/Cascade ecoregion north and west of the ecoregion divide, and in the Sierra Nevada south and east of the ecoregion divide.

### 2.2. Genetics Lab Work

We removed intestinal contents by squeezing them into sterile 10 mL tubes and then we dried the contents in a SpeedVac for 2-7 hours depending on the sample volume. We extracted DNA from two 150 mg subsamples of intestinal contents using an IBI Fecal DNA (IBI Scientific, Dubuque, IA, USA) extraction kit with bead beating for 120 seconds using bead beating tubes containing ceramic beads. We quantified DNA concentrations for all extractions using a qubit fluorometer (Invitrogen, Waltham, MA, USA).

We developed primers for this study targeting ∼150-200 bp fragments of mtDNA from each of the major taxonomic groups: frog, snake, lizard, mammal, salamander, insect, arachnid, gastropod, annelid, fish, and bird. Our custom primers were designed based on available mtDNA genomes and gene fragments for possible prey species downloaded from GenBank (Clark et al. 2016), including native, introduced, and domestic species in the Pacific Northwest. Detailed primer development methods are found in Appendix 1, and details of the primers and blocking primers for this study are presented in Table A2.

We amplified a fragment of the 12S, 16S, 18S, cytochrome C oxidase I (COI), or D-Loop gene in prey DNA using single primer PCR reactions for five primers (insects, annelids, fish, owls, and general birds) and multiplexed PCR reactions for 11 primers (lizard and snake, arachnid and gastropod, salamander and frog, mammal and Columbiformes, and Anatidae, Falconiformes/Accipiterformes, and Galliformes). We tagged the forward and reverse primers with an Illumina TruSeq Universal adapter to allow for the attachment of unique barcodes. For PCR master mixes and parameters, see Table A3, Table A4, and Table A5. We cleaned amplified PCR product with AMPure XP (Beckman Coulter, Brea, CA, USA) beads and quantified DNA concentrations of cleaned amplified product using a Qubit fluorometer. We combined the cleaned, amplified PCR products for each DNA extract into two libraries. The first library (blocking library) consisted of those primer sets that had blocking primers and the second library (non-blocking library) consisted of those primers that did not have blocking primers. We split our amplified products into two libraries to allow for higher depth of sequencing for those primers that amplified both host and prey DNA. We combined the amplified products for each DNA extract equimolar, then custom i5 and i7 indices with stem adapters were attached using KAPA HiFi HotStart ReadyMix (KAPA Biosystems). We cleaned the indexed products with AMPure XP beads to remove any unincorporated indices, then quantified concentrations and combined all samples equimolar into each library. Our libraries consisted of 80 samples including three duplicates, one owl positive control, one prey positive control that consisted of positive amplified products for each primer set, and a negative control. We sequenced the non-blocking library on Illumina MiSeq 2×250 nano runs with one library per lane and the blocking library on Illumina MiSeq 2×150 micro runs with one library per lane at the University of Wisconsin-Madison Biotechnology Center.

We used QIIME2 (Bolyen et al. 2019) to filter and process raw sequence data for the blocking library and non-blocking library separately. First, we used DADA2 (Callahan et al. 2016) plug-in to trim and filter our demultiplexed sequences and identify unique reads for each DNA sample. We built two custom sequence databases (blocking primers and non-blocking primers) with possible prey sequences downloaded from GenBank (Clark et al., 2016) and the sequenced prey from our primer optimization (Table A6). In each database, we also included barred owl and spotted owl sequences for the identification of host sequences. To filter out spurious sequences before taxonomic assignment, we excluded sequences that did not query against our database based on BLAST search. We trained a naïve Bayes classifier on our custom database in QIIME2 and then assigned molecular operational taxonomic units (MOTUs) to our sequenced reads. We re-ran this procedure with multiple filtering, trimming, and taxonomic assignment cut-offs to maximize the number of identified taxa and minimize spurious or incorrectly classified MOTUs. To further ensure correct taxonomic assignment, we performed closed and opened clustering. We BLAST searched (Altschul et al., 1990) any taxonomic assignment with less than 95% confidence against the NCBI database (Sayers et al., 2022) to confirm taxonomic assignment and percent sequence coverage. We then further filtered our final assignments by removing any assignment to an individual sample with less than 30 reads for both libraries. Finally, we merged the filtered read-count and MOTUs tables from both libraries for downstream analyses.

### 2.3. Estimating the occurrence of prey types

We determined the presence/absence of MOTUs for each barred owl sample and organized MOTUs into resource matrices, and estimated measures of prey occurrence for the taxonomic ranks of class, order, family, genus, and species. We also estimated occurrence metrics across these same ranks including only the vertebrate prey types to limit observations to the typically higher-biomass items. We estimated two measures of occurrence to describe the consumption of different prey types: (*i*) frequency of occurrence (FOO), or the percentage of barred owls in which each prey type was detected, and (*ii*) weighted percent of occurrence (wPOO), or the percentage of total diet each prey type represented with each sample contributing equally. After estimating occurrence measures, we established four datasets representative of barred owl diet for further analyses. These representative datasets, reflecting prey occurrence at the family and species level, and either including all prey types or only including vertebrate prey types, were designated as “All.Family”, “Verts.Family”, “All.Species”, and “Verts.Species”. Prey types with less than 1% wPOO within the representations were excluded from these datasets to minimize rarely utilized resources from affecting downstream inferences (Deagle et al., 2018).

### 2.4. Estimating the Diversity of Prey Types Consumed by Barred Owls

To test for sufficient sampling for diversity measures, we extrapolated our complete prey data at the lowest determinable taxonomic ranks through Hill numbers of order q=0 for species richness and q=1 for Shannon’s Diversity Index (Alberdi and Gilbert, 2019) using the package *iNEXT* (Hsieh et al., 2020) in program R version 4.0.2 (R Core Team, 2020). We compared the alpha diversity of prey types among the following categories: age class (adult or subadult), sex (male or female), and ecoregion (Klamath/Cascade or Sierra Nevada) for each of the four barred owl datasets using Shannon’s Diversity Index and the R package *vegan* (Oksanen et al., 2020). To determine the “average” barred owl diet within subgroups, we rarefied occurrence matrices for each diet representation and estimated the significance of distances between group centroids using Raup-Crick beta-diversity distances and the *vegan* functions “vegdist” to compute dissimilarity (variance) indices, “betadisper” to test for homogeneity of variances, and the permanova function “adonis2” (McArdle and Anderson, 2001) with 999 permutations to determine if estimated centroids differed between groups. To further compare the community of prey types in barred owl diets between ages, sexes, and ecoregions, we estimated beta-diversity for each category using Sørensen’s Index and the R package *betapart* (Baselga and Orme, 2012), and a one-way ANOVA to determine the significance of differences in variation among individuals within each category. We selected Sørensen’s Index so that the measure could be separated into each of its two components — turnover (i.e, species replacement) and nestedness (i.e., species gain/loss) — and their comparative contribution to the total observed variance estimated. In the case that prey use was not homogenous between subgroups, we used Tukey’s HSD test to determine which group within a category demonstrated greater variation. Additionally, we rarefied our data using the R package *phyloseq* (McMurdie and Holmes, 2013) and performed the same diversity tests to observe the effects of sample size on our results.

### 2.5 Effects of Environmental Features on Barred Owl Prey Consumption

To assess how prey consumption was shaped by environmental features, we characterized the landscape by vegetation conditions, aquatic features, and topography within barred owl home ranges. Home ranges were approximated by a 2,523m radius buffer centered on the removal location, approximating the 2,000ha home range size of GPS tagged barred owls in the Sierra Nevada ecoregion (Wood et al., 2020). We characterized vegetation using 2017 gradient nearest neighbor (GNN) data, which interpolates information from an extensive forest-inventory plot network across the landscape (30 m resolution) using Landsat satellite imagery (Ohmann and Gregory, 2002). We divided areas of the landscape into four classes — open, young, medium, and mature forest — based canopy cover and the quadratic mean diameter (QMD) of dominant and codominant trees corresponding to previous owl studies in the region (e.g. Hobart et al. 2019; Table A7). We then calculated the amount of each habitat class, as well as the basal area of hardwoods, within each home range (Table A7). As a proxy for the potential availability of aquatic prey within barred owl home ranges, we calculated the sum of the lengths of aboveground waterways (National Hydrography Dataset; https://www.usgs.gov/core-science-systems/ngp/national-hydrography/national-hydrography-dataset) and the length of shoreline of larger water bodies (see: https://hub.arcgis.com/datasets/esri::usa-detailed-water-bodies/about; Table A7). Finally, we calculated the mean elevation within each barred owl home range using digital elevation models (DEM) with a resolution of approximately 30m (USGS, 2018) (Table A7).

To estimate how prey communities varied as a function of environmental features, we performed constrained ordination using a redundancy analysis (RDA), where significance was tested through ANOVA-like permutations (n=999 permutations) and environmental variables were fit to the ordination through the permutation function “envfit” (n=99 permutations), all through the R-package *vegan* (Oksanen et al., 2020). Relationships between individual prey types and environmental features were assessed using logistic regression, where the binomial response variable reflected the presence (1) or absence (0) of the prey item. We fit generalized linear models of prey presence for all classes with a wPOO of ≥ 1% at its lowest determinable taxon, using all 124 sampled barred owls.

## 3. Results

### 3.1. Genetic Analyses

We sequenced 129 intestinal DNA extracts from lethally collected barred owls. The mean number of reads for the blocking library and non-blocking library was 56,161 and 15,270, respectively. Both library’s mean quality score was 37, indicating sufficient read coverage and quality. Two juveniles were removed from subsequent analyses as they are not considered independent dietary samples. Of the 127 remaining extracts, three were removed after filtering for read count for a final sample size of 124 (Table A1; Table A8).

We identified 78 MOTUs to a unique prey type at their lowest determinable taxonomic rank, which represented 10 different taxonomic classes. The majority of MOTUs (67.9%) were identified to the species rank. Of the remaining MOTUs, 23.1% could be identified to genus, and 7.7% to family. Only one prey type, representing an unknown spider in the order Araneae, could not be determined to the family rank or lower (Table A9).

### 3.2. Occurrence of Prey Types

Vertebrate prey accounted for 61.5% of the 78 identified prey types and was present in 80.6% of the 124 sampled owls (FOO), representing all major vertebrate groups of mammals, birds, reptiles, amphibians, and fish. Mammals accounted for the majority of vertebrate prey by FOO (64.5%) followed by the classes Amphibia (32.3%), Aves (21.8%), Reptilia (19.4%), and Actinopterygii (0.8%; Table A9). When considering only vertebrate prey types and wPOO, mammals again accounted for the majority of prey (52.8%), followed similarly by the classes Amphibia (21.4%), Aves (13.9%), Reptilia (11.4%), and Actinopterygii (0.5%; Fig. 2; Table A10). At lowest determinable taxons, northern flying squirrel (*Glaucomys sabrinus*) was the most common vertebrate prey type, occurring in 21.8% of sampled barred owls. Douglas squirrel (*Tamiasciurus douglassi*) and Pacific tree frog (*Pseudacris regilla*) represented the next most common vertebrate prey types. Additional small mammalian and amphibian prey, such as brush mouse (*Peromyscus boylii*), dusky-footed woodrat (*Neotoma fuscipes*), toads (*Anaxyrus* spp.), and Sierra tree frogs (*Pseudacris sierra*) were also common, with representatives of the Aves and Reptilia classes occurring less frequently. We detected a single incidence of fish predation (rainbow trout; *Onocorhynchus mykiss*). Barred owls also predated domestic species, such as domestic chickens (*Gallus gallus*) and cats (*Felis catus*), as well as game species such as ruffed grouse (*Bonansa umbellus*) and sooty grouse (*Dendragapus fuliginosus*; Table 1).

**Fig. 2.**
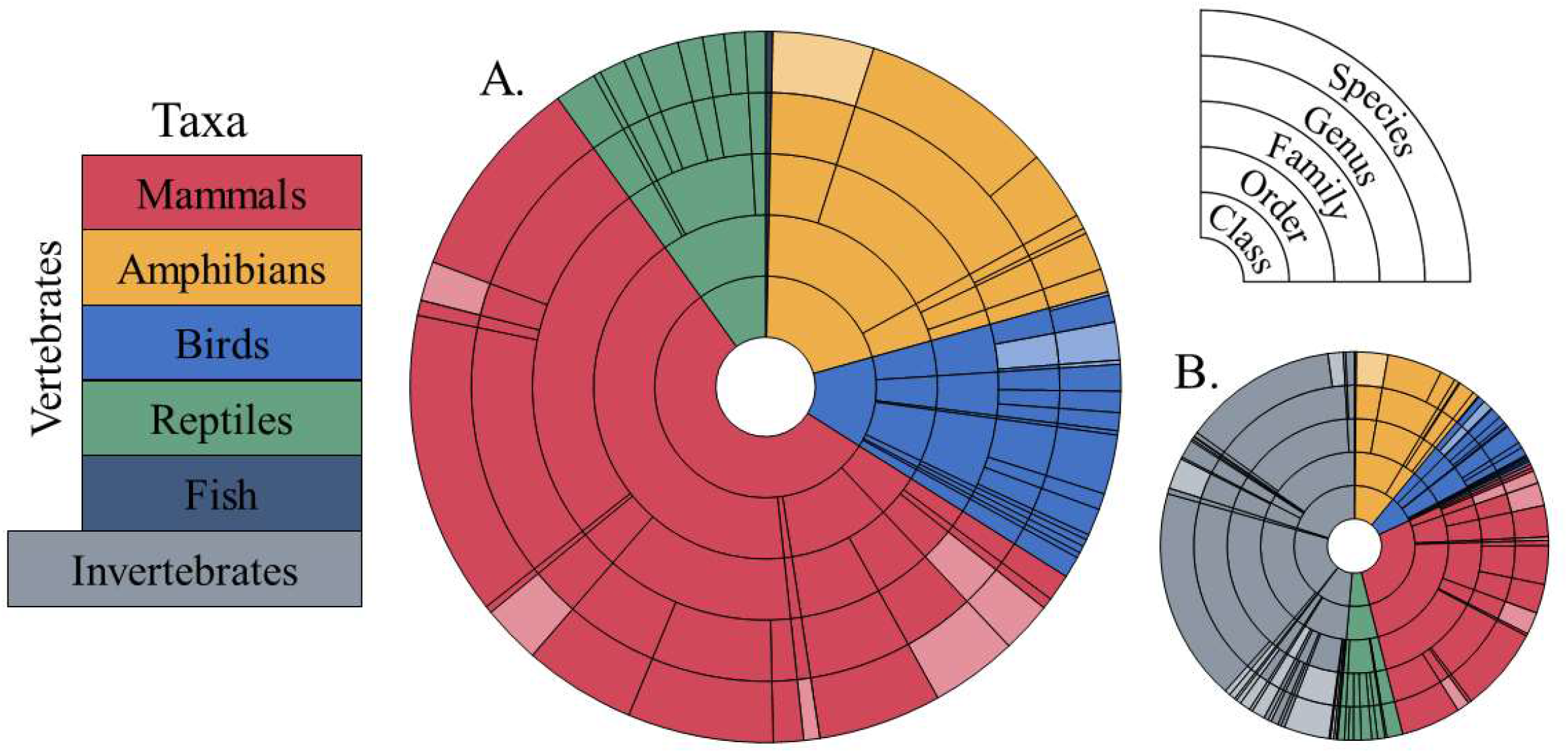
Proportion of prey types consumed by barred owls across taxonomic rank for (**A**) only vertebrate prey types and (**B**) both invertebrate and vertebrate prey types by weighted Percent of Occurrence (wPOO). Faded slices represent prey types that could not be determined beyond a higher taxonomic rank. Note that fish occurred at less than 1% wPOO across all rankings.

**Table 1.**
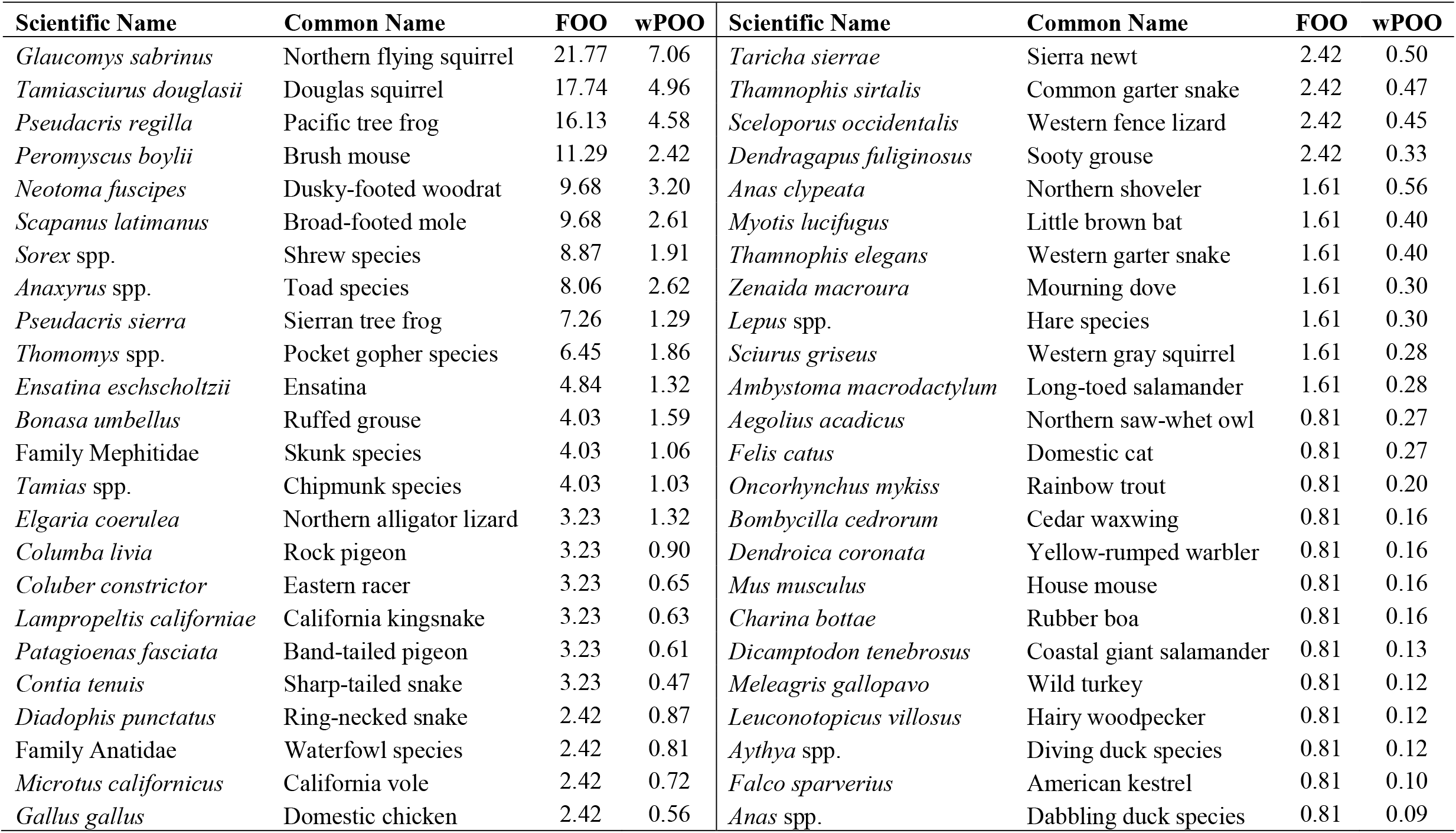
Vertebrate prey types’ frequency of occurrence (FOO) and weighted percent of occurrence (wPOO) at lowest determinable taxonomic level.

Vertebrate and invertebrate prey occurrences were evenly represented in the diet, with vertebrates accounting for a combined 51.1% wPOO at the family taxonomic level, and 51.4% wPOO based on all MOTUs identified to their lowest determinable rank (Fig. 2). Invertebrate prey types were detected across five classes: Arachnida, Chilopoda, Clitellata, Diplopoda, and Insecta. Common invertebrate prey included earthworms (Class: Clitellata), occurring in 51.6% of barred owl samples (FOO) at the class level, and Jerusalem crickets (*Stenopelmatus fuscus*), occurring in 41.1% of barred owls sampled (FOO) at the species level (Table A9).

We were unable to identify 53.3% of the 30 invertebrate prey types to the species level, resulting in the All.Species representative dataset being dropped from further analysis. The All.Family dataset was able to include all barred owls with successful extractions (n=124), while both the Verts.Family and Verts.Species datasets were constructed as to only include barred owls which had vertebrate prey types present after removing prey types with less than 1% wPOO (n=100, and n=99; Table A8).

### 3.3. Estimating the Diversity of Prey Types Consumed by Barred Owls

Extrapolation predicted a species richness of 108 (SE = 16), demonstrating insufficient sampling in our study for richness comparisons among groups of owls. Extrapolation of Shannon’s diversity index reached an asymptote within the interpolated samples however, suggesting sufficient sampling for estimating and testing diversity measures (Fig. A1). Barred owls collected from the Klamath/Cascade ecoregion demonstrated significantly greater alpha-diversity (mean Shannon’s Index = 1.02, SE = 0.05; *p* = 0.02) than those collected from the Sierra Nevada (mean = 0.77, SE = 0.05) based on the All.Family data representation. Barred owls in the Klamath/Cascade ecoregion also consumed a greater alpha-diversity of prey types (mean Shannon’s Index = 0.65. SE = 0.07) than those in the Sierra Nevada (mean = 0.42; SE = 0.08; *p* = 0.04) based on the Verts.Family data representation. Males and females consumed a similar diversity of prey across all three representative datasets, as did adults and subadults, with all *p*-values exceeding 0.4.

Barred owls consumed similar prey within all categories, with no significant differences observed between rarefied Raup-Crick centroids (minimum *p* = 0.43). Variation in prey consumption differed between ecoregions for barred owls in the Verts.Species dataset (mean Sørensen’s Index = 0.63, SE = 0.008; *F*_1,97_=5.78, *p*=0.018), with greater variation in diet among individuals in the Sierra Nevada than the Klamath/Cascade (Tukey HSD; *p* = 0.038, SE = 0.017). Both turnover (mean = 0.61, SE= 0.011; *F*_1,97_ = 4.41, *p* = 0.038) and nestedness (mean = 0.089, SE = 0.008, *F*_1,97_ = 9.19, *p* = 0.003) were significant contributors to the total beta-diversity, with more of the total beta-diversity explained by the turnover component (Fig. A2). There was not a significant difference in variation in either the sex or age categories, nor in any categories in the All.Family or Verts.Family datasets (minimum *p* = 0.08). These same tests, conducted with rarefied data, did not produce any significant results.

### 3.4. Effects of Environmental Features on Barred Owl Prey Consumption

Environmental features did not explain a significant amount of variation in prey communities for the All.Family (5.5%), Verts.Family (6.0%), and Verts.Species (6.4%) data representations based on redundancy analyses (minimum p-value = 0.59). Prey types with ≥1% wPOO at their lowest taxonomic rank consisted of 2 families, 6 genera, and 13 species. Elevation was a significant predictor of occurrence for two prey species based on logistic regressions. Barred owls were more likely to consume Pacific tree frogs with increasing elevation (β = 0.001, SE = 0.0007, *p* = 0.04), and more likely to consume broad-footed moles (*Scapanus latimanus*) with decreasing elevation (β = -0.04, SE = 0.002, *p* = 0.01). Additionally, Pacific tree frogs were less likely to be consumed within home ranges with a higher basal area of hardwoods (β = -0.13, SE = 0.06, *p* = 0.04). At the generic rank, longhorn beetles (*Monochamus* spp.) were consumed more often with increasing elevation (β = 0.002, SE = 0.0008, *p* = 0.03), and less likely to be consumed in home ranges with a high basal area of hardwoods (β = -0.17, SE = 0.08; *p* = 0.04) or water features (β = -0.07, SE = 0.03; *p* = 0.02).

## 4. Discussion

Previous dietary studies of barred owls in their invasive range have been limited to morphological examinations of prey remains in nests, pellets, and stomachs, and visual observations of feeding (Livezey, 2007; Wiens et al., 2014), and in one case stable isotopes in feathers (Wood et al., 2021), and thus may not capture the full suite of prey consumed. Our customized DNA barcoding approach demonstrated that barred owls have a broad diet in their invasive range, consuming numerous mammalian, amphibian, avian, reptilian, and invertebrate prey. Furthermore, barred owls consumed a higher proportion of non-mammalian prey than observed in previous studies in the invasive part of their range (Hamer et al., 2001; Wiens et al., 2014, Livezey, 2007; Fig. 2). Some of these differences were likely the result of geographic variation in prey communities, given the different areas in which these studies were performed. However, we suggest that differences in barred owl diet between studies also resulted from increased PCR sensitivity to prey types that would either be unlikely to form a pellet or visually absent due to rapid degradation. While our study was not designed to test for the effects of barred owl predation or competition on wildlife species and communities, the apparent generalist predatory strategy of barred owls — coupled with densities that can achieve 8-10x that of displaced spotted owls (Forsman et al., 2011; Wiens et al. 2011, 2014) — supports concerns over the potential widespread ecological impacts of this invasive species. Our comprehensive identification of prey species provides a basis for the further ecological studies needed to understand the population level impacts barred owls may have on native species, as well as effects on biological communities and ecosystems.

### 4.1. Barred owls as direct predators

Our results suggest that barred owls will not serve as ecological replacements for the spotted owls they displace. Spotted owls are highly specialized on small mammals, particularly flying squirrels and woodrats (Verner et al., 1992; Hamer et al., 2001; Wiens et al., 2014; Zulla et al., 2022). Invasive barred owls are likely to increase the predation pressure faced by these species, due to the barred owls’ larger body size and greater population density (Holm et al., 2016). Barred owls may also exert novel predation pressure on mammal species which are rare if not absent in the diets of spotted owls using traditional methodologies.

Beyond mammals, barred owls consumed a variety of prey taxa not typically observed in spotted owl diets when assessed from pellet analysis (Forsman et al., 1984; Smith et al., 1999) or from camera-based observations of nest deliveries (Zulla et al., 2022). Barred owl consumption of amphibians appears to be prevalent. Amphibian declines have historically been severe in western North America (Corn, 1994), and may be further exacerbated by the addition of a novel, avian predator. Avian prey was also detected at a higher rate than in spotted owl diets. Our finding of a northern saw-whet owl (*Aegolius acadius*) lends credence to concerns of intra-guild predation as a possible explanation regarding a loss of insular populations of other small forest owls in the Pacific Northwest following barred owl establishment (Acker, 2012). Finally, reptiles, although nearly absent from previous studies of barred owl diet in the Pacific Northwest (Hamer et al., 2001; Wiens et al., 2014), were prevalent in our study. Thus, our genetic based diet assessment points to potential direct effects of barred owls on both previously suspected prey and novel taxa not observed in prior studies, and at rates much higher than would be expected in native spotted owls.

While we did not detect any species currently listed as threatened or endangered under the federal or state Endangered Species Acts, such species are typically rare and incidental predation would likely be difficult to ascertain. This is particularly true because genetic screening of intestinal samples only assesses a small temporal snapshot of an individual’s broader diet. Yet, the prevalence of amphibian predation illustrates a likely risk to threatened and endangered species. For example, although our observed cases of toad predation were geographically isolated from known threatened species and populations, barred owls have been observed and collected from areas of overlap with the federally threatened Yosemite toad (*Anaxyrus canorous;* Hofstadter et al., 2022).

Invertebrates accounted for approximately half of detected prey types by wPOO, and likely constitute important prey resources for barred owls. Direct predation of the two most commonly detected invertebrate species, earthworms (*Lumbricus rubellus*) and Jerusalem crickets, was confirmed by observation of stomach contents (*personal observation*) and prey deliveries to nests (Elderkin, 1987). Our study was limited to presence/absence data however, and beyond these two highly prevalent invertebrate species we were thus unable to quantify number of predation events or total prey biomass — two metrics that may better explain the importance of small-bodied prey. We were also unable to distinguish between primary and possible secondary predation (prey consumption by species consumed by barred owls). Nonetheless, while our study was not fully optimized for the detection and assessment of invertebrate prey resources, we still demonstrated that some invertebrates are prevalent in western barred owl diets.

### 4.2. Barred owls as competitors of native predators

Our analysis also supports concerns that barred owls may compete with native predators for prey resources (Holm et al. 2016). Flying squirrels and dusky-footed woodrats constitute the primary prey of spotted owls in many regions (Verner et al., 1992), and these key spotted owl prey types constituted the first and third most common mammalian prey of barred owls sampled in this study. Furthermore, Douglas squirrels, while uncommon in the diet of spotted owls (e.g., Hamer et al., 2001), were the second most common vertebrate species consumed by barred owls and can be a key prey resource for sensitive predators in the region including the Pacific fisher (*Pekania pennanti*; Golightly et al., 2006) and northern goshawk (*Accipiter gentilis*; Keane et al., 2006). High densities of barred owls relative to spotted owls may be particularly facilitated by the inclusion of the Douglas squirrel in their diet, given that this prey species is typically large in body mass than flying squirrels (∼225g versus ∼166g; Smith, 2003), and likely occurs at similar densities (∼1acre^-1^; Smith, 1968; Williams et al., 1992). This may constitute a competitive advantage of barred owls over spotted owls, while also leading to further competition within native predator assemblages.

### 4.3. Factors shaping barred owl prey consumption

While barred owl prey consumption generally varied little based on intrinsic factors (i.e., sex and age), we observed significant variation in diet among the Klamath/Cascade and Sierra Nevada ecoregions based on measure of both alpha- and beta-diversity. We suggest that these differences are likely a product of either local or seasonal prey availability — two factors this study cannot assess. However, high-occurrence families were among the most evenly represented prey types between the two ecoregions (Fig. 3). These groups likely demonstrate the preferred prey of barred owls, and do not appear to be the drivers of differences in alpha- and beta-diversity. Differences in diversity metrics appear to have been driven by less frequently occurring prey groups that are more likely to be only incidentally consumed. Thus, geographic variation in the consumption of these less frequent families is likely the result of differences in prey abundance and availability.

**Fig. 3.**
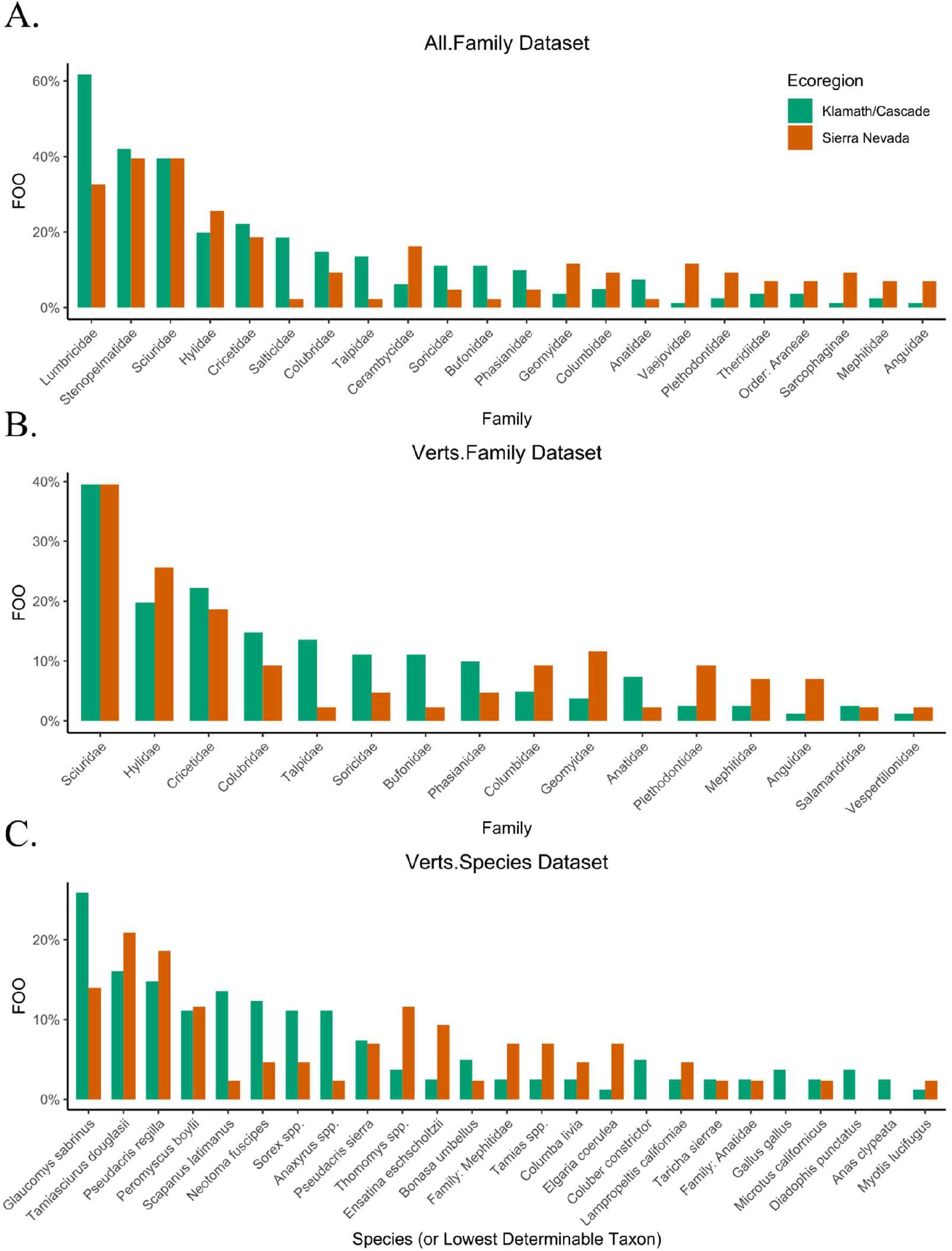
Differences in frequency of occurrence (FOO) rates of common (>1% weighted percent occurrence) prey species consumed by barred owls between the Sierra Nevada and Klamath/Cascade ecoregions, driving alpha- and beta-diversity results in the (A) All.Family, (B) Verts.Family, and (C) Verts.Species datasets.

At local scales approximating home ranges, we detected only six statistically significant relationships between prey type consumption and the seven environmental features considered. Elevation was the most likely to be biologically important and accounted for three of the significant relationships, likely driven by elevation gradients in prey distribution. Overall, however, the lack of strong relations between environmental characteristics and prey species again highlights the strong generalist tendencies of western barred owls.

## 5. Management Implications

Experimental studies conducted in the Pacific Northwest and Sierra Nevada have demonstrated that lethal removals can substantially reduce barred owl densities and alleviate well-documented competitive pressures on native spotted owls (Diller et al., 2014; Wiens et al., 2021; Hofstadter et al., 2022). Our results suggest that removals conducted to promote the recovery of spotted owls may also benefit a broad range of additional vertebrate species consumed by invasive barred owls. While barred owl populations have been reduced to very low densities throughout the range of the California spotted owl in the Sierra Nevada (Hofstadter et al., 2022), removal areas only encompass a small portion of the range of the northern spotted owl in California, Oregon, and Washington (Wiens et al., 2021). Without expanding current efforts, barred owl populations will inevitably continue to grow outside of the limited footprints of these areas. Further, several removal studies in the range of the northern spotted owl have ended, and these areas will likely be rapidly recolonized by barred owls from surrounding areas (Van Lanen et al., 2011; Diller et al., 2014; Wiens et al., 2021). We recommend that managers and decision makers continue to adhere to the Precautionary Principle of early and rapid management for barred owls in areas where the invasion is relatively recent (Wood et al., 2020). This approach would involve continued removals in the range of the California spotted owls, as well as additional proactive removals in the few remaining areas of low barred owl densities within the range of the northern spotted owl. Such proactive efforts could prevent potential effects on biological communities before they occur, requiring less effort and resources to curb barred owl populations than in areas where populations have become established (Diller et al., 2014; Hofstadter et al. 2022). Removals in areas where barred owl populations have achieved high densities, however, will require substantially more effort and resources, but may be needed to reverse impacts to native species. Ecological studies conducted in conjunction with barred owl removals would further help elucidate the effects of these abundant apex predators on biological communities in the Pacific Northwest.

Here, we demonstrate that lethal management methods, in conjunction with developing technologies such as seen here with DNA metabarcoding and high-throughput-sequencing, can provide information critical to assessing community-wide effects of invasive predators. We uncovered numerous affected prey types that were not previously detected with traditional methodologies, and the absence of this information could have negatively impacted the ability of managers to appropriately respond to the invasive species. We recommend that when the Precautionary Principle is applied to invasive predators, consideration should be given to not just the immediate effects of removing predator species, but to the utility of such methods in understanding the full scope of community effects.

## Acknowledgements

We thank WJ Berigan, BP Dirksmeyer, and MM Barker for assisting with barred owl removals, and A Reiss and the many acoustic and field technicians who conducted surveys for barred owls. We also thank C Cicero, TLW Barclay, S Medina, M Paramonova, and additional Museum of Vertebrate Zoology students and staff for assistance in dissecting barred owl specimens. We thank C Kulzer, the California Academy of Sciences, the Museum of Vertebrate Biology, and additional researchers for the collection and loaning of prey reference samples.

## Abbreviations

MOTU: Molecular Operational Taxonomic Unit
FOO: Frequency of Occurrence
wPOO: Weighted Percent of Occurrence
QMD: Quadratic Mean Diameter
DEM: Digital Elevation Model
GLM: Generalized Linear Model
RDA: Redundancy Analysis
GNN: Gradient Nearest Neighbor
CDFW: California Department of Fish and Wildlife
CNDDB: California Natural Diversity Database

