## Supplemental Materials for "DNA metabarcoding reveals the threat of rapidly expanding barred owl populations to native wildlife in western North America"

**Supplementary Materials**

### A1. Detailed Primer Development Methods

#### A1.1 Primer/Blocking Primer Creation

We developed a list of potential prey that included species found in redwood forests in California as well as species found in old growth forests in the Pacific Northwest from wildlife occurrence records (gbif.org; inaturalist.org; wildlife.ca.gov; spiderid.com; usaspiders.com; insectidentification.org; Roth and Sadeghian, 2006; United States Bureau of Land Management, 1999; Reynolds, 2017). For primer creation, potential prey species were separated into major taxonomic groups (i.e., frog, snake, lizard, mammal, salamander, insect, arachnid, gastropod, annelid, fish, and bird). To allow for more accurate species assignments, bird species were further separated into taxonomic family groups for Anatidae, Columbiformes, Falconiformes, Accipiterformes, and Galliformes.

We obtained mitochondrial genomes from potential prey species from GenBank (Clark et al. 2016). If no mitochondrial genome was available for the target prey species, we obtained mitochondrial genomes from congener species, if available. We imported the mitochondrial genomes into MEGA-X (Kumar et al., 2018) and aligned the sequences using the CLUSTAL W algorithm (Thompson et al., 1994). Using the Primaclade (Gadberry et al. 2005) web-based platform (The Santos Lab http://webhome.auburn.edu/~santosr/primaclade.htm), we obtained a list of potential primers for each taxonomic group alignment. From this list, we discarded any primer pair that amplified a fragment length other than 150-200 base pairs (bp), had more than two degeneracies, or had repeated base pairs at the 3' end. To ensure no amplification of predator DNA, we then imported the mitochondrial genomes for *Strix varia* and *S. occidentalis occidentalis,* aligned these sequences with our groups of interest, and searched for the oligo motif in the predator sequences. If the oligo had a mismatch of four or fewer base pairs compared to the predator sequence, we determined if blocking primers could be created to prevent the amplification of predator DNA. If it was not possible to create blocking primers from any of the possible primer sets, we still retained the primers for downstream filtering.

To test for specificity in amplification of prey DNA or predator DNA outside of the mitochondrial genome, we used NCBI Primer BLAST (Ye et al. 2012). First, we blast searched the primers against the northern spotted owl (*S. occidentalis caurina*) reference genome (StrOccCau_2.0; Hanna et al., 2017). If primers amplified any fragment less than 1000 bp in the northern spotted owl genome, we discarded those primers. Next, we blast searched the primers against Family groupings of the potential prey and downloaded any sequence fragments that were found for our potential prey. We imported these sequences into MEGA-X and aligned them using the CLUSTAL W algorithm. We then searched for parsimony-informative sites between species to determine if our primers were able to distinguish between different species. Finally, we checked any remaining primers for hairpins and primer dimers using NEB Oligo Analyzer (Owczarzy et al. 2008). After all filtering stages, we created blocking primers following the methods of Vestheim and Jarman (2008) for any of the remaining primer sets. If it was not possible to create blocking primers, we kept the top five oligo sets in a taxa group for testing *ex situ*.

**A1.2 Positive controls for primer testing and reference database**

To test the efficacy of our primers, we obtained bones, feathers, scale clips or tissues from potential prey species or conspecifics of potential prey items (Table A6). We extracted DNA from feathers, scale clips, and tissue using a Qiagen DNeasy Blood and Tissue kit with an added step of bead beating homogenization for two minutes for hard tissue samples. To extract DNA from bones, we first crushed the bones to a powder using a mortar and then extracted DNA using a QIAmp Investigator kit.

We PCR amplified genomic DNA using the primers designed for each taxonomic group. Each PCR reaction consisted of 1X PCR buffer, 400 μM of dNTPs, 0.8 μM of forward and reverse primer, 0.5 mM $MgCl_2$, 0.6 mg/mL BSA, 1 unit taq polymerase (Phusion, New England Biolabs), 2 μL DNA, and water to total 25 μL. For those taxonomic groups that we created blocking primers for, each PCR reaction also consisted of 9 μM of forward and reverse blocking primers. We included a PCR negative in each run consisting of water. Barred owl and California spotted owl DNA was tested with each primer set to ensure that host DNA was not amplified. If a primer set failed to amplify any DNA for either prey or hosts during testing, we then tested the primer set without using blocking primers to ensure that the blocking primers were not blocking amplification of prey DNA. We checked for amplification of DNA and sequenced any positive amplification. Prior to sequencing, we cleaned all amplified products using ExoSAP IT (Affymetrix). We pair-end sequenced the fragments on an ABI 3730xl DNA Analyzer at the Biotechnology Center, University of Wisconsin-Madison. GenBank accession numbers are provided in Table A6. We visualized the sequence chromatograms and aligned the forward and reverse sequences in MEGA-X. We BLAST (Altschul et al., 1990) searched the sequences to determine identity. All sequences were included in the reference database for assigning molecular taxonomic units.

#### A1.3 Primer optimization

After finding the top primer set for each taxonomic group, we then tested if primer sets could be multiplexed in PCR reactions using the program MFEprimer (Wang et al., 2019). To test the multiplex reactions, we mixed DNA from different taxonomic groups into a single reaction. We first tested the multiplex with Phusion taq and, if Phusion taq failed to amplify any samples, we then tested Qiagen UCP multiplex taq. We tagged all our primers with the Illumina TruSeq Universal adapter and once again tested the primers for specificity in amplifying only prey DNA and to optimize the annealing temperature.

#### A.1 References


Altschul, S. F., Gish, W., Miller, W., Myers, E. W., & Lipman, D. J. (1990). Basic local alignment search tool. *Journal of molecular biology*, *215*(3), 403-410. https://doi.org/10.1016/S0022-2836(05)80360-2

California Department of Fish and Wildlife. https://wildlife.ca.gov/ (accessed May 2019)

Clark, K., Karsch-Mizrachi, I., Lipman, D. J., Ostell, J., & Sayers, E. W. (2016). GenBank. *Nucleic acids research*, *44*(D1), D67-D72. https://doi.org/10.1093/nar/gkv1276

Gadberry, M. D., Malcomber, S. T., Doust, A. N., & Kellogg, E. A. (2005). Primaclade—a flexible tool to find conserved PCR primers across multiple species. *Bioinformatics*, *21*(7), 1263-1264. https://doi.org/10.1093/bioinformatics/bti134

GBIF | Global Diversity Information Facility. http://www.gbif.org/ (accessed May 2019)

Hanna, Z.R., Henderson, J.B., Wall, J.D., Emerling, C.A., Fuchs, J., Runckel, C., Mindell, D.P., Bowie, R.C., DeRisi, J.L. and Dumbacher, J.P., 2017. Northern spotted owl (Strix occidentalis caurina) genome: divergence with the barred owl (Strix varia) and characterization of light-associated genes. *Genome biology and evolution*, *9*(10), 2522-2545.

iNaturalist. http://www.inaturalist.org/ (accessed May 2019)

Insect Identification. https://www.insectidentification.org/ (accessed May 2019)

Kumar, S., Stecher, G., Li, M., Knyaz, C., & Tamura, K. (2018). MEGA X: molecular evolutionary genetics analysis across computing platforms. *Molecular biology and evolution*, *35*(6), 1547. https://doi.org/10.1093/molbev/msy096

Owczarzy, R., Tataurov, A. V., Wu, Y., Manthey, J. A., McQuisten, K. A., Almabrazi, H. G., Pedersen, K. F., Lin, Y., Garretson, J., McEntaggart, N. O., Sailor, C. A., Dawson, R. B., & Peek, A. S. (2008). IDT SciTools: a suite for analysis and design of nucleic acid oligomers. *Nucleic acids research*, *36*, W163–W169. https://doi.org/10.1093/nar/gkn198

Reynolds, J.W., 2017. A summary of the status of earthworms (Annelida: Oligochaeta) in ecoregions of the United States.

Roth, B. and Sadeghian, P.S., 2006. *Checklist of the land snails and slugs of California* (p. 81). Santa Barbara (CA): Santa Barbara Museum of Natural History.

Spider ID. https://spiderid.com/ (accessed May 2019)

Thompson, J. D., Higgins, D. G., & Gibson, T. J. (1994). CLUSTAL W: improving the sensitivity of progressive multiple sequence alignment through sequence weighting, position-specific gap penalties and weight matrix choice. *Nucleic acids research*, *22*(22), 4673-4680. https://doi.org/10.1093/nar/22.22.4673

United States. Bureau of Land Management. Oregon State Office, 1999. *Field guide to survey and manage terrestrial mollusk species from the Northwest Forest Plan*. Bureau of Land Management, Oregon State Office.

USA Spiders. Spiders in California. https://usaspiders.com/spiders-in-california/ (accessed May 2019)

Vestheim, H. and Jarman, S.N., 2008. Blocking primers to enhance PCR amplification of rare sequences in mixed samples–a case study on prey DNA in Antarctic krill stomachs. *Frontiers in zoology*, *5*(1), 1-11.

Wang, K., Li, H., Xu, Y., Shao, Q., Yi, J., Wang, R., Cai, W., Hang, X., Zhang, C., Cai, H., & Qu, W. (2019). MFEprimer-3.0: quality control for PCR primers. *Nucleic acids research*, *47*(W1), W610-W613. https://doi.org/10.1093/nar/gkz351

Ye, J., Coulouris, G., Zaretskaya, I., Cutcutache, I., Rozen, S., & Madden, T. L. (2012). Primer-BLAST: a tool to design target-specific primers for polymerase chain reaction. *BMC bioinformatics*, *13*(1), 1-11. https://doi.org/10.1186/1471-2105-13-13

**Table A1.** Sample and corresponding Museum of Vertebrate Zoology catalogue IDs

| Sample ID | Catalogue ID |
|---|---|
| 19 | MVZ:192320 |
| 21 | MVZ:192327 |
| 32 | MVZ:192329 |
| 33 | MVZ:192309 |
| 34 | MVZ:192344 |
| 40 | MVZ:192336 |
| 44 | MVZ:192311 |
| 57 | In preparation |
| 61 | In preparation |
| 65 | In preparation |
| 73 | MVZ:193282 |
| 90 | In preparation |
| 149 | MVZ:193283 |
| 150 | In preparation |
| 151 | MVZ:193267 |
| 152 | In preparation |
| 685 | MVZ:192343 |
| 686 | MVZ:192332 |
| 703 | MVZ:192323 |
| 722 | MVZ:192346 |
| 724 | MVZ:192318 |
| 740 | MVZ:192352 |
| 742 | MVZ:192321 |
| 753 | MVZ:192315 |

| | |
|---|---|
| 754 | MVZ:192333 |
| 755 | MVZ:192328 |
| 756 | MVZ:192345 |
| 778 | MVZ:192306 |
| 779 | MVZ:192319 |
| 786 | MVZ:192310 |
| 787 | MVZ:192312 |
| 790 | MVZ:192334 |
| 793 | MVZ:192322 |
| 796 | MVZ:192317 |
| 797 | MVZ:192314 |
| 806 | MVZ:192340 |
| 810 | MVZ:192326 |
| 811 | MVZ:192313 |
| 812 | MVZ:192307 |
| 816 | MVZ:192338 |
| 817 | MVZ:192341 |
| 818 | MVZ:192308 |
| 819 | MVZ:192330 |
| 820 | MVZ:192342 |
| 835 | MVZ:193278 |
| 837 | In preparation |
| 838 | In preparation |
| 839 | In preparation |
| 841 | MVZ:193260 |
| 842 | In preparation |
| 843 | In preparation |

| | |
|---|---|
| 845 | In preparation |
| 846 | In preparation |
| 848 | In preparation |
| 849 | In preparation |
| 850 | MVZ:193264 |
| 851 | In preparation |
| 852 | In preparation |
| 853 | In preparation |
| 854 | In preparation |
| 862 | MVZ:193247 |
| 864 | In preparation |
| 865 | In preparation |
| 866 | In preparation |
| 867 | In preparation |
| 868 | In preparation |
| 869 | In preparation |
| 871 | In preparation |
| 873 | In preparation |
| 874 | In preparation |
| 875 | In preparation |
| 877 | In preparation |
| 902 | MVZ:193249 |
| 903 | MVZ:193262 |
| 904 | MVZ:193250 |
| 905 | MVZ:193259 |
| 906 | In preparation |
| 907 | In preparation |

| | |
|---|---|
| 909 | In preparation |
| 910 | MVZ:193288 |
| 922 | MVZ:193245 |
| 923 | MVZ:193265 |
| 924 | MVZ:193255 |
| 925 | MVZ:193277 |
| 926 | MVZ:193252 |
| 929 | MVZ:193251 |
| 930 | MVZ:193248 |
| 931 | MVZ:193284 |
| 932 | MVZ:193253 |
| 933 | MVZ:193257 |
| 934 | MVZ:193270 |
| 935 | MVZ:193287 |
| 936 | MVZ:193290 |
| 937 | MVZ:193256 |
| 938 | MVZ:193258 |
| 939 | MVZ:193272 |
| 940 | MVZ:193269 |
| 941 | In preparation |
| 942 | In preparation |
| 943 | In preparation |
| 944 | In preparation |
| 945 | In preparation |
| 946 | MVZ:193286 |
| 947 | In preparation |
| 948 | In preparation |

| | |
|---|---|
| 949 | MVZ:193281 |
| 950 | MVZ:193279 |
| 951 | MVZ:193274 |
| 952 | MVZ:193261 |
| 954 | In preparation |
| 955 | MVZ:193289 |
| 957 | MVZ:193276 |
| 981 | In preparation |
| 982 | MVZ:193246 |
| 983 | MVZ:193271 |
| 984 | MVZ:193273 |
| 986 | MVZ:193280 |
| 987 | MVZ:193292 |
| 988 | MVZ:193291 |
| 989 | MVZ:193285 |
| 990 | MVZ:193263 |
| 991 | MVZ:193254 |
| 992 | MVZ:193266 |
| 993 | In preparation |

**Table A2.** The primers and blocking primers (BP) used this study to amplify prey DNA from barred owl intestinal contents. All primers were created for this study except the fish 16S primers which were from Deagle et al. 2007. Primers were tagged with an Illumina adapter to allow for the attachment of individual barcodes. Taq polymerases used in this study were: Phusion High-Fidelity DNA Polymerase (New England Biolabs, cat M0530), UCP Multiplex PCR (Qiagen, cat 206742), and AllTaq Master Mix (Qiagen, 203144).

| Primer Name | Primer Sequence | Gene Fragment | Annealing Temperature (°C) | Taq Polymerase |
|---|---|---|---|---|
| Lizard For. | GGTATYCTAACCGTGCAAAG | 16S | 63 | Phusion High-Fidelity |
| Lizard Rev. | GAAGTCGCCCCAACTYAA | | | |
| Lizard BP For. | GCAATCAATTGTCTCATAAATCGG | | | |
| Lizard BP Rev. | CGAAAAAGTGGCGGTCAA | | | |
| Snake For. | CCAGAAGACCCTGTGAAGC | | | |
| Snake Rev. | GCGCTGTTATCCCTGGAGTA | | | |
| Snake BP For. | GTGGAACTTAAAAATCACCGGTC | | | |
| Snake BP Rev. | GGTAGCTTGGTCCATTGTTCAAT | | | |
| Arachnid For. | ACGAGCGCTTTTATTAGACC | 18S | 62 | Phusion High-Fidelity |
| Arachnid Rev. | ATGTGGTAGCCGTTTCTCAG | | | |
| Gastropod For. | GCTGCWACTATARTTATTGSAGTACC | COI | | |
| Gastropod Rev. | CNGCAAAWAGRGCAAATACAGC | | | |

| | | | | |
|---|---|---|---|---|
| FalcoAccip For. | GRAGTGGAAGWAATGGGCTACA | 12S | 60 | Phusion High-Fidelity |
| FalcoAccip Rev. | ACGACTTACCTCATCTTYGGC | | | |
| FalcoAccip BP For. | GCTACACTCTCTGYCARCAGA | | | |
| FalcoAccip BP Rev. | TAGCCTGGGTGGTGRTATAGG | | | |
| Galliformes For. | CAACCCCTGCCTRTAATGTAC | D-Loop | | |
| Galliformes Rev. | AGARGATGCCGCGATCAC | | | |
| Anatidae For. | GCACCTAAACACACCATYAAGATGATC | 12S | | |
| Anatidae Rev. | GCACCGCCAAGTCCTTAGAG | | | |
| Anatidae BP For. | AAACGACCCYAGCCCTGC | | | |
| Anatidae BP Rev. | CTGGCGGATGTTCTAGTGGTGATG | | | |
| Salamander For. | ATAAGACGAGAAGASCCTRTGGAGC | 16S | 60 | UCP Multiplex |
| Salamander Rev. | TTATCCCTRGSGTAACTTRGTTCG | | | |
| Salamander BP For. | AACTTAAAAATCACCSGTCACCCCG | | | |
| Salamander BP Rev. | AATTGGTATTGGATCTGGTTGCTG | | | |
| Frog For. | TATAAGACGAGAAGACSCYATGGA | | | |
| Frog Rev. | TGTTATCSCYAGGGTAASTTGGTTC | | | |

| | | | | |
|---|---|---|---|---|
| Frog BP For. | CTSTGGAACTTAAAAATCACCGG | | | |
| Frog BP Rev. | TTGTTSAATTGGTATTGGATCTGGTTGC | | | |
| Mammal For. | AAGACGAGAAGASCCTATGGAGC | 16S | 63 | UCP Multiplex |
| Mammal Rev. | TGCGCTGTTATCCCTAGSGTAA | | | |
| Mammal BP For. | GTGGAACTTAAAAATCACCGGTCACC | | | |
| Mammal BP Rev. | AGCTTGGTCCATTGTTCAATTGGTA | | | |
| Columbidae For. | TGTGGAACTTAAAAATCARCAGCC | | | |
| Columbidae Rev. | GGTAGCTTGGTCCATTGRTCA | | | |
| Columbidae BP For. | GTCACCCCGCACAAMCAC | | | |
| Columbidae BP Rev. | GAGGTGKTTGTGCGGGGTGA | | | |
| Insect For. | RGACGAGAAGACCCTATAGA | 16S | 56 | AllTaq |
| Insect Rev. | ACGCTGTTATCCCTAARGTAA | | | |
| Insect BP For. | GTGGAACTTAAAAATCACCGG | | | |
| Insect BP Rev. | GTAGCTTGGTCCATTGTTCA | | | |
| Annelid For. | ACAAGCTACCTTAGSGATAACAG | 16S | 62 | Phusion High-Fidelity |
| Annelid Rev. | TAGGTCCTTTCGTASTATAAATAGAT | | | |

| | | | | |
|---|---|---|---|---|
| Annelid BP For. | CCAAAGCTGSTATTCTMTACTAAAC | | | |
| Annelid BP Rev. | RTTGGATTTTAGTGGGGTTTCTC | | | |
| Fish 16S1F.[1] | GACGAKAAGACCCTA | 16S | 50 | Phusion High-Fidelity |
| Fish 16S2R.[1] | CGCTGTTATCCCTADRGTAACT | | | |
| GenBird For. | GTASCGTAAGGGAAAGRTGAAATA | 16S | 64 | Phusion High-Fidelity |
| GenBird Rev. | CACAGGYAACCAGCTATCAC | | | |

[1]Deagle, B. E., N. J. Gales, K. Evans, S. N. Jarman, S. Robinson, R. Trebilco, and M. A. Hindell (2007). Studying seabird diet through genetic analysis of faeces: a case study on macaroni penguins (*Eudyptes chrysolophus*). PLoS One 2:e831.

**Table A3.** The PCR master mix for all single-plex PCRs using Phusion High-Fidelity taq and AllTaq. All single-plex PCRs were conducted with a 25 µL total reaction volume with 2.5 µL DNA extract in each reaction.

| Reagent | Without Blocking Primers<br>Final Concentration | With Blocking Primers<br>Final Concentration |
|---|---|---|
| **Phusion High-Fidelity** | | |
| Water | 12.95 µL | 8.45 µL |
| 5X buffer | 1X | 1X |
| dNTPS (10 mM) | 400 µM | 400 µM |
| Forward primer (20 µM) | 0.8 µM | 0.8 µM |
| Reverse primer (20 µM) | 0.8 µM | 0.8 µM |
| Forward blocking primer (100 µM) | - | 9 µM |
| Reverse blocking primer (100 µM) | - | 9 µM |
| $MgCl_2$ (25 mM) | 0.5 mM | 0.5 mM |
| BSA (20 mg/mL) | 0.6 mg/mL | 0.6 mg/mL |
| Taq polymerase | 1.25 units | 1.25 units |
| **Phusion High-Fidelity (GenBird Primer Set Only)** | | |
| Water | 13.80 µL | - |
| 5X buffer | 1X | - |
| dNTPS (10 mM) | 400 µM | - |
| Forward primer (20 µM) | 0.8 µM | - |
| Reverse primer (20 µM) | 0.8 µM | - |
| Forward blocking primer (100 µM) | - | - |
| Reverse blocking primer (100 µM) | - | - |
| $MgCl_2$ (25 mM) | 0.2 mM | - |
| BSA (20 mg/mL) | 0.16 mg/mL | - |

| | | |
|---|---|---|
| Taq polymerase | 1.25 units | - |
| **AllTaq** | | |
| Water | - | 8.5 μL |
| AllTaq Master Mix | - | 6.25 μL |
| Forward primer (20 μM) | - | 0.8 μM |
| Reverse primer (20 μM) | - | 0.8 μM |
| Forward blocking primer (100 μM) | - | 9 μM |
| Reverse blocking primer (100 μM) | - | 9 μM |
| $MgCl_2$ (25 mM) | - | 0.5 mM |
| BSA (20 mg/mL) | - | 0.6 mg/mL |

**Table A4.** The PCR master mix for all multi-plex PCRs using Phusion High-Fidelity taq and UCP Multiplex. All multiplex PCRs using Phusion taq were conducted with a 50 µL total reaction volume with 5.0 µL DNA extract in each reaction. All multiplex PCRs using UCP were conducted with a 40 µL total reaction volume with 4.5 µL DNA extract in each reaction.

| | Two primer sets without blocking primers | Two primer sets with blocking primers | Three primer sets with blocking primers |
|---|---|---|---|
| Reagent | Final Concentration | Final Concentration | Final Concentration |
| **Phusion High-Fidelity** | | | |
| Water | 20.9 µL | 8.9 µL | 4.9 µL |
| 5X buffer | 1X | 1X | 1X |
| dNTPS (10 mM) | 400 µM | 400 µM | 400 µM |
| Forward primer (20 µM) | 0.8 µM | 0.8 µM | 0.8 µM |
| Reverse primer (20 µM) | 0.8 µM | 0.8 µM | 0.8 µM |
| Forward primer (20 µM) | 0.8 µM | 0.8 µM | 0.8 µM |
| Reverse primer (20 µM) | 0.8 µM | 0.8 µM | 0.8 µM |
| Forward primer (20 µM) | - | - | 0.8 µM |
| Reverse primer (20 µM) | - | - | 0.8 µM |
| Forward blocking primer (200 µM) | - | 12.0 µM | 12.0 µM |
| Reverse blocking primer (200 µM) | - | 12.0 µM | 12.0 µM |
| Forward blocking primer (200 µM) | - | 12.0 µM | 12.0 µM |
| Reverse blocking primer (200 µM) | - | 12.0 µM | 12.0 µM |

| | | | |
|---|---|---|---|
| Forward blocking primer (200 µM) | - | - | 12.0 µM |
| Reverse blocking primer (200 µM) | - | - | 12.0 µM |
| $MgCl_2$ (25 mM) | 0.5 mM | 0.5 mM | 0.5 mM |
| BSA (20 mg/mL) | 0.6 mg/mL | 0.6 mg/mL | 0.6 mg/mL |
| Taq polymerase | 1.25 units | 1.25 units | 1.25 units |
| **UCP (Frog/Salamander primer sets only)** | | | |
| Water | - | 8.0 µL | - |
| UCP Multiplex Master Mix | - | 10.0 µL | - |
| Forward primer (20 µM) | - | 0.8 µM | - |
| Reverse primer (20 µM) | - | 0.8 µM | - |
| Forward primer (20 µM) | - | 0.8 µM | - |
| Reverse primer (20 µM) | - | 0.8 µM | - |
| Forward blocking primer (200 µM) | - | 12.0 µM | - |
| Reverse blocking primer (200 µM) | - | 12.0 µM | - |
| Forward blocking primer (200 µM) | - | 12.0 µM | - |
| Reverse blocking primer (200 µM) | - | 12.0 µM | - |
| $MgCl_2$ (25 mM) | - | 0.31 mM | - |
| BSA (20 mg/mL) | - | 0.5 mg/mL | - |

| | | | |
|---|---|---|---|
| **UCP (Mammal/Columbidae primer sets only)** | | | |
| Water | - | 8.5 µL | - |
| UCP Multiplex Master Mix | - | 10.0 µL | - |
| Forward primer (20 µM) | - | 0.8 µM | - |
| Reverse primer (20 µM) | - | 0.8 µM | - |
| Forward primer (20 µM) | - | 0.8 µM | - |
| Reverse primer (20 µM) | - | 0.8 µM | - |
| Forward blocking primer (200 µM) | - | 12.0 µM | - |
| Reverse blocking primer (200 µM) | - | 12.0 µM | - |
| Forward blocking primer (200 µM) | - | 12.0 µM | - |
| Reverse blocking primer (200 µM) | - | 12.0 µM | - |
| $MgCl_2$ (25 mM) | - | 0.16 mM | - |
| BSA (20 mg/mL) | - | 0.33 mg/mL | - |

**Table A5.** The PCR parameters for all primer sets and taqs used in this study. All PCRs were run on a BioRAD S1000.

| | Temperature (°C) | Time | Number of cycles |
|---|---|---|---|
| **Phusion High-Fidelity: Single-plex** | | | |
| Initial Denaturation | 98 | 30 sec | |
| Denaturation | 98 | 20 sec | |
| Annealing | 50-64 | 30 sec | 40 |
| Elongation | 72 | 30 sec | |
| Final Elongation | 72 | 10 min | |
| **AllTaq** | | | |
| Initial Denaturation | 95 | 2 min | |
| Denaturation | 95 | 15 sec | |
| Annealing | 56 | 20 sec | 40 |
| Elongation | 72 | 20 sec | |
| Final Elongation | 72 | 5 min | |
| **Phusion High-Fidelity: Multiplex** | | | |
| Initial Denaturation | 98 | 30 sec | |
| Denaturation | 98 | 20 sec | |
| Annealing | 60-63 | 30 sec | 40 |
| Elongation | 68 | 30 sec | |
| Final Elongation | 68 | 10 min | |
| **UCP** | | | |
| Initial Denaturation | 95 | 2 min | |
| Denaturation | 95 | 15 sec | |
| Annealing | 60-63 | 30 sec | 40 |
| Elongation | 72 | 30 sec | |

| | | |
|---|---|---|
| Final Elongation | 72 | 5 min |

**Table A7.** Environmental variables, definitions, and data sources within 2,523m radius buffer centered on removal locations. Mature, medium, and young forests were determined by a combination of canopy cover and quadratic mean diameter (QMD) of standing trees. Data is sourced from 2017 gradient nearest neighbor (GNN), digital elevation model (DEM) value, and the National Hydrography Dataset (NHD) with additional information from the Environmental Systems Research Institute (ESRI).

| **Metric** | **Definition** | **Data Source** |
|---|---|---|
| Mature Forest | Proportional area with QMD >= 60cm, canopy cover >=40% | GNN |
| Medium Forest | Proportional area with QMD 30-60cm, canopy cover >=40% | GNN |
| Young Forest | Proportional area with QMD < 30cm, canopy cover >=40% | GNN |
| Open Forest | Proportional area with Canopy cover <40% | GNN |
| Hardwoods | Mean basal area of hardwoods in territory | GNN |
| Elevation | Mean elevation | DEM |
| Water Features | Summed lengths of available shoreline and water channels | NHD and ESRI |

**Table A8:** Sample counts for categorical variables in each representative dataset. Samples of unknown age were not included in age comparisons.

| | | Representative Dataset | | |
|---|---|---|---|---|
| | | All.Family | Verts.Family | Verts.Species |
| Age | Adult | 97 | 78 | 78 |
| | Subadult | 24 | 19 | 18 |
| | Unknown | 3 | 3 | 3 |
| Sex | Male | 62 | 47 | 47 |
| | Female | 62 | 53 | 52 |
| Ecoregion | Klamath | 81 | 65 | 64 |
| | Sierra Nevada | 43 | 35 | 35 |
| | Total | 124 | 100 | 99 |

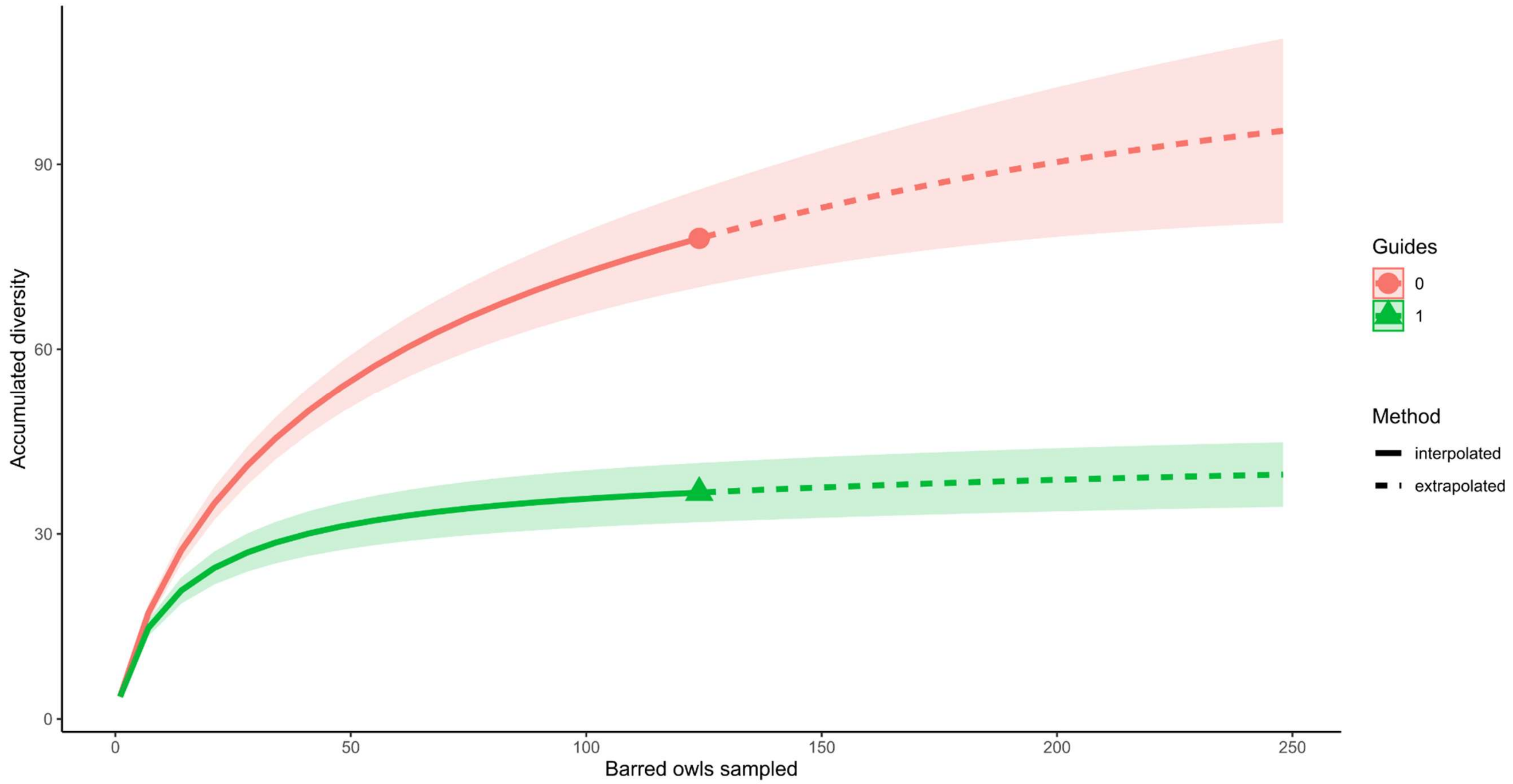


**Fig. A1.** Interpolated and extrapolated diversity by Hill numbers of order q=0 (species richness) and q=1 (Shannon's entropic index), demonstrating insufficient sampling for species richness but sufficient sampling for Shannon's index.

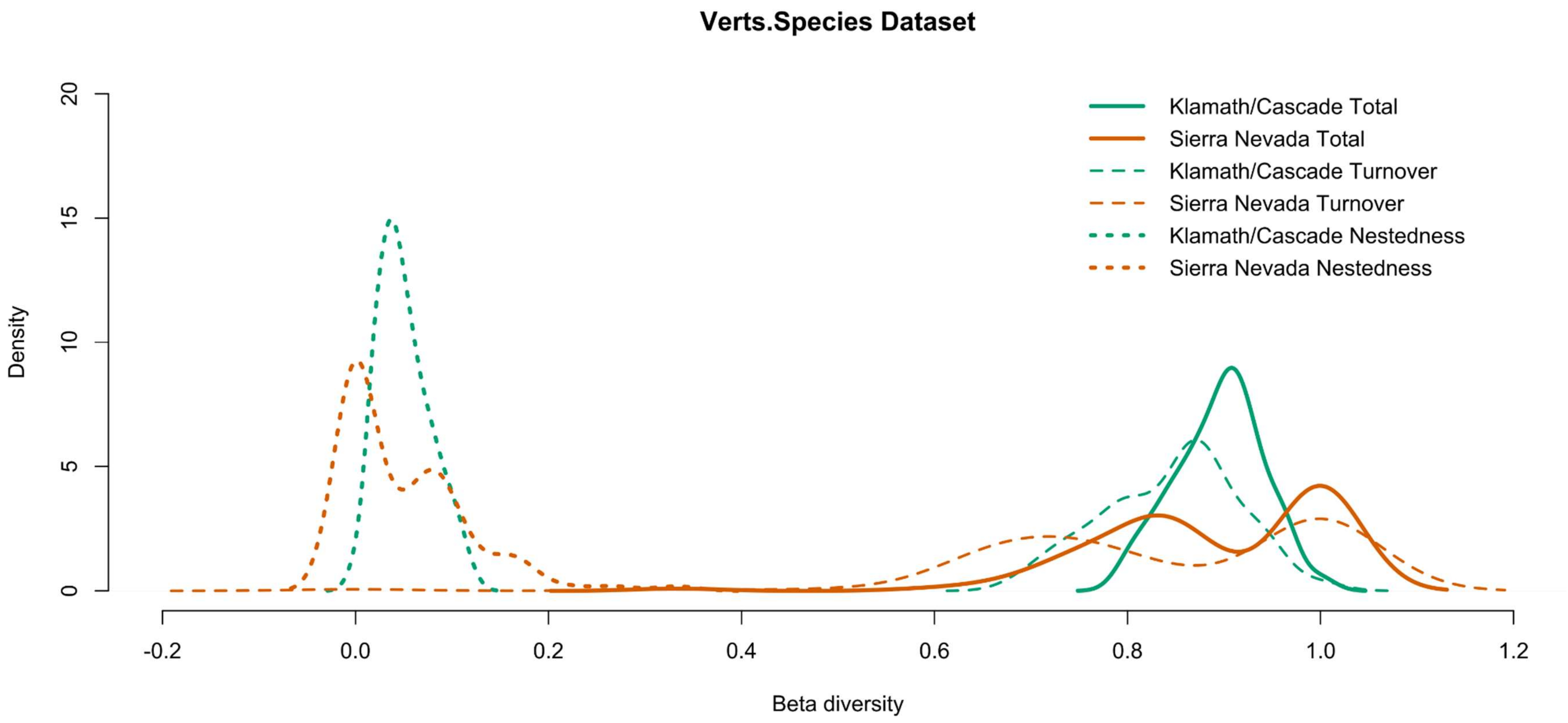


**Fig. A2.** Density of beta-diversity totals, and both components of turnover and nestedness, between the Sierra Nevada and Klamath/Cascade ecoregions in the Verts.Species dataset of barred owl prey types.
